# Anaerobic methane oxidizing archaea offset sediment methane concentrations in Arctic thermokarst lagoons

**DOI:** 10.1101/2022.06.20.496783

**Authors:** Sizhong Yang, Sara E. Anthony, Maren Jenrich, Michiel H. In ‘t Zandt, Jens Strauss, Pier Paul Overduin, Guido Grosse, Michael Angelopoulos, Boris K. Biskaborn, Mikhail N. Grigoriev, Dirk Wagner, Christian Knoblauch, Andrea Jaeschke, Janet Rethemeyer, Jens Kallmeyer, Susanne Liebner

## Abstract

Thermokarst lagoons represent the transition state from a freshwater lacustrine to a marine environment, and receive little attention regarding their role for greenhouse gas production and release in Arctic permafrost landscapes. We studied the fate of methane (CH_4_) in sediments of a thermokarst lagoon in comparison to two thermokarst lakes on the Bykovsky Peninsula in northeastern Siberia through the analysis of sediment CH_4_ concentrations and isotopic signature, methane-cycling microbial taxa, sediment geochemistry, and lipid biomarkers. We specifically assessed whether sulfate-driven anaerobic methane oxidation (S-AOM) through anaerobic methanotrophic archaea (ANMEs), common in marine sediments with constant supply of sulfate and methane, establish after thermokarst lagoon development and whether sulfate-driven ANMEs consequently oxidize CH_4_ that would be emitted to the water column under thermokarst lake conditions. The marine-influenced lagoon environment had fundamentally different methane-cycling microbial communities and metabolic pathways compared to the freshwater lakes, suggesting a substantial reshaping of microbial and carbon dynamics during lagoon formation. Anaerobic sulfate-reducing ANME-2a/2b methanotrophs dominated the sulfate-rich sediments of the lagoon despite its known seasonal alternation between brackish and freshwater inflow. CH_4_ concentrations in the freshwater-influenced sediments averaged 1.34±0.98 µmol g^−1^, with highly depleted δ^13^C-CH_4_ values ranging from -89‰ to -70‰. In contrast, the sulfate-affected upper 300 cm of the lagoon exhibited low average CH_4_ concentrations of 0.011±0.005 µmol g^−1^ with comparatively enriched δ^13^C-CH_4_ values of -54‰ to -37‰ pointing to substantial methane oxidation. Non-competitive methylotrophic methanogens dominated the methanogenic community of the lakes and the lagoon, independent of porewater chemistry and depth. This potentially contributed to the high CH_4_ concentrations observed in all sulfate-poor sediments. Our study shows that S-AOM in lagoon sediments can effectively reduce sediment CH_4_ concentrations and we conclude that thermokarst lake to lagoon transitions have the potential to mitigate terrestrial methane fluxes before thermokarst lakes fully transition to a marine environment.

## Introduction

Northern permafrost contains an estimated 1,035 ± 150 PG of organic carbon within the top three meters (Hugelius et al. 2014), which is more than 30% of total terrestrial soil organic carbon (Strauss et al. 2021) and disproportionally high given its limited areal coverage of 15 - 22% of the terrestrial area (Schuur et al. 2015). The warming climate has triggered permafrost degradation with a variety of consequences for the landscape (Biskaborn et al. 2019), including top-down thaw (increasing active layer) over long term, thermo-erosion along coasts, rivers, and lake shores, rapid thawing associated with thermokarst processes (Strauss et al. 2017; Turetsky et al. 2020) and significant land subsidence (Anders et al. 2020; Antonova et al. 2018). Thermokarst develops in the lowland areas of ice-rich permafrost, and typically results in formation of thermokarst ponds or lakes (Bouchard et al. 2016). Thermokarst landscapes are estimated to cover approximately 20-40% of permafrost regions (Olefeldt et al. 2016; Strauss et al. 2017). Thermokarst lakes are hotspots of CH_4_ emission (Walter Anthony et al. 2016), especially those lakes that formed in organic-rich Yedoma permafrost, which contains approximately 40% of total permafrost carbon (Strauss et al. 2017; Zimov et al. 2006). As CH_4_ has approximately 28 times the global warming potential of carbon dioxide (CO_2_) over 100 years (Dean et al. 2018; Myhre et al. 2013), excessive CH_4_ emissions from thermokarst landscapes can yield disproportionate CO_2_ equivalents and feedback to the climate system (Serikova et al. 2019; Walter Anthony et al. 2018).

The oceans only contribute 1-2% of global CH_4_ emissions even though they cover approximately 70% of the earth’s surface, due to effective reduction of CH_4_ emissions through anaerobic methane oxidation coupled with sulfate reduction termed S-AOM (Hinrichs and Boetius 2002; Knittel et al. 2019; Wallenius et al. 2021). S-AOM consortia between anaerobic methanotrophic archaea (ANMEs) and sulfate-reducing bacteria are widespread in marine systems, although ANMEs are very slow growing (Knittel and Boetius 2009). Thermokarst lagoons represent a transitional state between freshwater thermokarst lakes and a fully marine environment. Lagoon formation occurs when intrusion of marine water introduces high concentrations of ions, especially sulfate, to the geochemical profiles of previous freshwater lakes (Spangenberg et al. 2021). Vertical diffusion of these salts also changes the ion composition of the freshwater sediments, creating a salt gradient across several meters of sediment (Angelopoulos et al. 2020). The sediment geochemistry of thermokarst lagoons largely depend on their hydrological connection with the sea. For example, open lagoons have a constant exchange with the sea leading to a more stable mixing regime and higher salinity and sulfate concentrations, while semi-closed lagoons have a seasonal connection that is often interrupted during winter months due to ice thickening resulting in seasonally more variable environmental conditions (Jenrich et al. 2021; Schirrmeister et al. 2018).

The role of thermokarst lagoons for carbon turnover and greenhouse gas release have received little attention so far despite their role in coastal carbon dynamics (in ‘t Zandt et al. 2020) and widespread occurrence along Arctic coasts (Angelopoulos et al. 2021). Marine-water inundated systems may have the potential to effectively reduce CH_4_ emissions through establishment of sulfate-reducing anaerobic methane oxidation (S-AOM). Because ANME organisms are ubiquitous in marine environments where CH_4_ and sulfate are present, they potentially also establish in Arctic lagoons. However, there is currently very little literature on the microbial consortia in freshwater systems inundated with marine water. One of the few examples comes from Weil et al. (2020), a study which included the ‘Karrendorfer Wiesen’, a formerly drained peatland re-wetted with marine water in 1993, which has no established S-AOM community even more than 25 years after the rewetting. In a rewetted coastal fen near Rostock in Germany with infrequent marine water intrusion events, only archaea from the so called ANME-2d group were found which are known to use nitrate as electron acceptor but again no sulfate-driven ANMEs (like ANME 2a/2b) were observed (Wen et al. 2018). One study in tropical coastal lagoons by Chuang et al. (2017) found very little methane oxidation activity even though sulfate and CH_4_ concentrations were high. For Arctic coastal lagoons, a pilot study on the CH_4_ concentrations in winter ice cores recently provided evidence that CH_4_ is oxidized in brackish water beneath the ice cover during progressive ice formation in the winter (Spangenberg et al. 2021) but studies on the fate of CH_4_ in Arctic lagoon sediments are missing. Unlike the poor evidence for S-AOM in transitionary habitats between land and sea, ANME 2a/2b were found to thrive in deep submarine permafrost (Winkel et al. 2018), indicating that permafrost deposits in a fully marine environment are suitable for supporting S-AOM.

We hypothesize that Arctic thermokarst lagoons could be suitable, albeit non-canonical, environments for the establishment of sulfate-driven anaerobic methane oxidizers, and that these organisms could consequently oxidize large amounts of CH_4_ in comparison to freshwater thermokarst lake sediments. We investigated sediments of two thermokarst lakes and a thermokarst lagoon on the Bykovsky Peninsula in northeastern Siberia and analyzed sediment CH_4_ concentrations and isotopic signature, methane-cycling microbial taxa, sediment geochemistry, and lipid biomarkers to understand how the formation of thermokarst lagoons influences microbial CH_4_ formation and oxidation.

## Material and Methods

### Site Description

The study area is located on the Bykovsky Peninsula, southeast of the Lena Delta in the Buor-Khaya Gulf of the Laptev Sea in northeastern Siberia, Russia (Fig 1). Three of the water bodies on the peninsula were selected for this study: two freshwater thermokarst lakes, Lake Golzovoye (also anglicized as Goltsovoye, abbreviated as LG in this study) and Northern Polar Fox Lake (LNPF) and one thermokarst lagoon, Polar Fox Lagoon (PFL). Lake Golzovoye is relatively young, and formed approximately 8000 years cal BP (Jongejans et al. 2020). Northern Polar Fox Lake and Polar Fox Lagoon lie within the same partially drained thermokarst basin (Angelopoulos et al. 2020), with the lake to the north of the lagoon. The basin may have gone through repeated draining events, and initial thermokarst lake formation occurred between 12.5 and 9.4 cal ka BP (Grosse et al. 2007; Jenrich 2020). The lagoon is a nearly-closed system with a wide, shallow and winding channel supplying the lake with water from Tiksi Bay during the summer, with large seasonal variation in salinity and ion concentrations such as sulfate and chloride (Angelopoulos et al. 2020; Spangenberg et al. 2021).

**Fig. 1.**
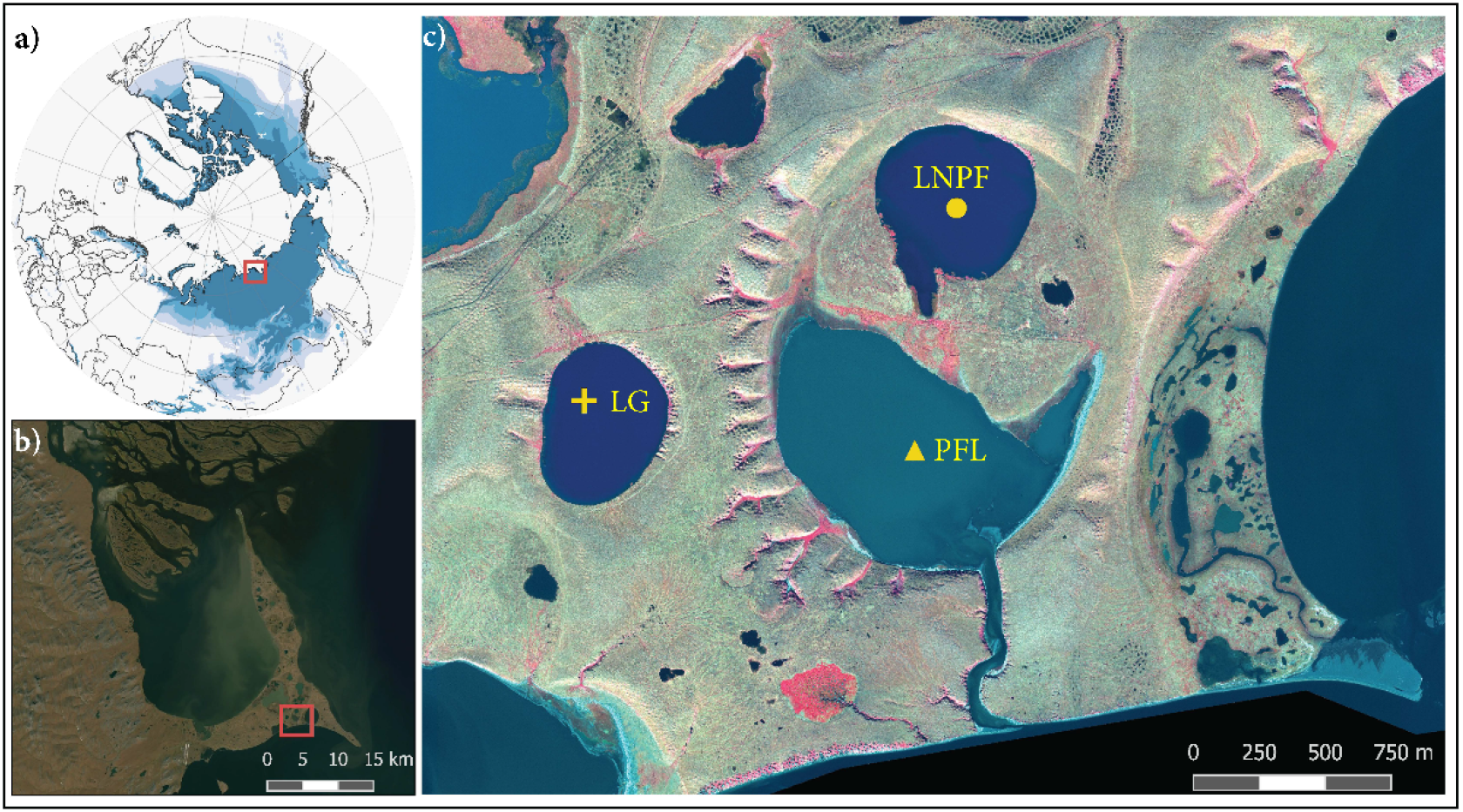
Location and map of the study site showing a) location with respect to the Northern Hemisphere and permafrost extent regions b) location with respect to the Bykovsky Peninsula, and c) relative location of the lakes and the lagoon. Note the small channel at the bottom of PFL connecting to Tiksi Bay. Symbols show location of coring sites: Triangle - Polar Fox Lagoon; Circle - Northern Polar Fox Lake; Cross - Lake Golzovoye. Map in a) based on Brown et al. (1997), b) ESRI Satellite World Imagery, and c) based on WorldView-3 false color satellite image (8-5-3), acquired 2016-09-02 (©DigitaGlobe).

### Field Work and Sampling

During a drilling campaign on the Bykovsky Peninsula in April 2017, cores PG2420, PG2426, and PG2423 were retrieved from LG, LNPF, and PFL, with drilling depths from the sediment surface of 5.2, 5.4, and 6.1 m, respectively. Water height above sediment surface was 510, 560, and 310 cm for LG, LNPF and PFL, respectively. The cores were recovered with a hammer-driven 60 mm Niederreiter piston corer (UWITEC™) in overlapping sections retrieved in 3-m-core barrels. All cores consisted of unfrozen sediment at time of sampling. After retrieval, the cores were cut into 10 cm liner segments. The segments were packed in N_2_-flushed, vacuum sealed opaque bags and were stored at approximately 4°C (pore water analysis) or frozen (molecular analysis, lipid biomarkers, bulk analysis) before processing. Sampling for sediment CH_4_ concentrations and isotopes as well as molecular work was done in a cleaned and weather-protected tent at the coring site without time delay after a core was retrieved. Defined sediment plugs (3 cm^3^) were obtained from the edges of the 10 cm core slices using cut syringes and placed into 20 ml vials containing saturated NaCl (gas analysis) or into sterile 15 ml falcon tubes (molecular work). A total of 139 of 10cm segments were used for analyses: 41 from LG, 39 from LNPF, and 59 from PFL.

### Porewater sampling and analysis

Porewater samples were extracted using a hydraulic press in an anoxic glovebox and then filtered to 0.45 µm microporous membrane. pH was measured with a WTW MultiLab 540 probe (WTW, Germany). Samples were analyzed by suppressed ion chromatography for all major cations and anions. Alkalinity was measured by colorimetric titration. Dissolved iron (ferric and ferrous) concentrations in pore water were measured via spectrophotometry by the ferrozine method (Viollier et al. 2000), with a detection limit of 0.25 μM. Samples were measured at GFZ German Research Centre for Geosciences.

### CH_4_ concentrations and carbon isotopes

Concentrations of CH_4_ were determined by static headspace gas chromatography (GC7890, Agilent Technologies, USA) with a flame ionizing detector (FID). Plug samples were mixed into a slurry before analysis, and concentrations are presented in units of micromoles of CH_4_ per gram of whole sediment plug (µmol g^−1^). Stable carbon isotope signatures of CH_4_ were analyzed by isotope-ratio mass spectrometry (Delta V plus, Thermo Scientific, Dreieich, Germany) coupled to a GC-Isolink / Trace-GC (Thermo Scientific, Dreieich, Germany) at the Universität Hamburg. Values are expressed relative to Vienna Peedee belemnite (VPDB) using IAEA NGS3, (-73.3‰ VPDB) as external standards. The analytical error of these analyses was ± 0.2 ‰.

### Bulk Parameters

Samples were analyzed for total carbon (TC), total organic carbon (TOC), total nitrogen (TN), and total sulfur (TS) at the University of Cologne. For analysis of TC, TN, and TS, 20-30 mg aliquots of dry sample were combined with 20 mg of tungsten trioxide (an oxidation catalyst) and were measured on an Elementar Micro Vario elemental analyzer. For TOC measurements, samples were first processed to remove inorganic carbon by adding 1% HCl to 20-50 mg of dry bulk sample for 1 h at 60 °C, and then allowed to continue to react overnight at room temperature. Samples were then thoroughly rinsed with 18.2 MΩ purified water until pH was back to neutral values and dried in the oven at 60°C. Afterwards, 20 mg of tungsten oxide were added to each and samples were analyzed using the same Elementar Micro Vario elemental analyzer.

### Lipid Biomarker Indices

Lipid extraction was performed to obtain *n*-alkanes for use in several biomarker indices to characterize the origin of organic matter (OM) in the lakes. Samples (6 - 10 g dry weight) were ultrasonically extracted following the method of Bligh and Dyer (1959). The total lipid extracts were then separated into neutral lipid, glycolipid and phospholipid fractions using hand-packed SiOH columns and eluting with dichloromethane, acetone, and methanol, respectively. The neutral fractions (containing *n*-alkanes) were further separated using activated SiO_2_ column chromatography and *n*-hexane as eluent. The *n*-alkanes were measured on a gas chromatograph equipped with an on-column injector and flame ionization detector (GC-FID, 5890 series II, Hewlett Packard, USA). A fused silica capillary column (Agilent DB-5MS; 50 m × 0.2 mm; film thickness: 0.33 µm) was used with helium as carrier gas. *n-Alkane* identification and quantification was performed using an external standard mixture of C_21_-C_40_ *n-alkanes* (PN 04071, Sigma-Aldrich).

Detailed information about the indices used can be found in the corresponding references, but briefly: The carbon preference index (CPI) indicates the maturity of the OM, where higher values are generally indicative of younger, less degraded OM. The CPI is calculated after Marzi et al. (1993):

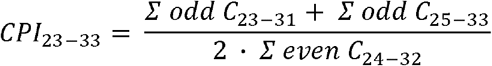

The proxy P_aq_ was proposed by Ficken et al. (2000) and first applied to permafrost by Zheng et al. (2007). The P_aq_ expresses the portion of OM derived from submerged or floating macrophytes (P_aq_ > 0.4) as opposed to emergent and terrestrial plant input (P_aq_ < 0.4). We use the general trend of P_aq_ rather than the absolute values due to the proxy being applied to permafrost regions only recently, and not developed for this soil type.

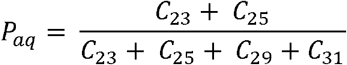

### DNA extraction and Illumina sequencing of amplicons and metagenomic DNA

Total nucleic acids were extracted in duplicate using the PowerSoil-Kit (MO-BIO, Qiagen) according to the manufacturer’s protocol. The concentration of DNA extracts was checked via gel electrophoresis and on a Qubit fluorometer (Invitrogen).

Amplicon libraries were prepared with the barcoded primer pair Uni515-F/Uni806-R to cover both bacterial and archaeal 16S rRNA genes (Caporaso et al. 2011). Each sample was run in technical duplicates. The 50 µL PCR reactions contained 10x Pol Buffer C (Roboklon GmbH, Berlin, Germany), 25 mM MgCl_2_, 0.2 mM deoxynucleoside triphosphate (dNTP) mix (ThermoFisher Scientific), 0.5 mM each primer (TIB Molbiol, Berlin, Germany) and 1.25 U of Optitaq Polymerase (Roboklon). The PCR program started with a denaturation step at 95 °C for 10 min, followed by 35 cycles at 95 °C for 15 s, annealing at 60 °C for 30 s, extension at 72 °C for 45 s and a final extension step at 72 °C for 5 min. The tagged PCR products were then purified with the Agencourt AMPure XP kit (Beckman Coulter, Switzerland) and eluted in 30 µL DNA/RNA-free water. The purified product was quantified and then the equilibrated PCR, together with positive and negative controls, were pooled. The sequencing was performed on an Illumina MiSeq system (paired-end, 2 × 300 bp) by Eurofins Scientific (Konstanz, Germany).

In addition to amplicon sequencing, metagenomic sequencing was specifically done on six samples of sediment from PFL by using the Illumina HiSeq 2500 platform by Eurofins Scientific (Konstanz, Germany). These six samples are representative of the sulfate zone (combination of 40, 100, and 130 cm), the upper (190 cm), middle (220 cm), and bottom (250 cm) of the Sulfate-Methane transition zone, freshwater sediment (combination of 310, 420, and 480 cm), and permafrost sediment (570 cm).

### Data processing

For amplicon data, the raw sequences were processed according to Yang et al. (2021a). The community composition was reported in ASV (amplicon sequence variants) tables and the taxonomy was assigned against the SILVA138 database (https://www.arb-silva.de). In this study, special attention was paid to methanogens and aerobic/anaerobic methanotrophic oxidizers. The ASV table generated by QIIME2 was filtered by taxonomy to keep only methane producing archaea, and methanotrophic oxidizers. The duplicates of each sample were merged. The resulting data set has 264 ASVs including methanogenic archaea (‘Meth’ hereafter), aerobic methane oxidizing bacteria (MOB) and anaerobic oxidizers of methane (AOM) over 23 samples.

Metagenomic data for PFL were processed with steps of quality control, assembly, genomic binning, and functional annotation according to the flow described by Yang et al. (2021b). The abundance of key functional genes (in Transcripts Per Kilobase Million, TPM) involved in methanogenesis, aerobic methane oxidation (MOB), dissimilatory nitrate reduction (DNR) and sulfate reduction (DSR) was compared to provide additional insight to the amplicon-derived data for PFL. To check the taxonomy bearing these genes, the translated protein sequences of these genes were taxonomically annotated by DIAMOND (v0.8.22.84) (Buchfink et al. 2015) against UniProt90 reference database (https://www.uniprot.org).

### Network analysis

First, the ASVs with mean relative abundances less than 0.05% across all samples were removed from downstream analysis. The resultant dataset was further filtered to keep those ASVs with a Spearman correlation coefficient > 0.6 and statistically significant p value < 0.01 with custom scripts in R (version 4.0.2) (R_Core_Team 2014). These filtering steps removed poorly represented ASVs and reduced network complexity, facilitating the determination of the non-random co-occurrence of the core community. The network was then explored and visualized using the *igraph* package (v 1.2.5) (Csardi and Nepusz 2006). Community modularity was detected via a *walktrap* algorithm according to the internal ties and the pattern of ties between different groups with the *igraph* package.

The association between the members (ASVs) of each module and environment variables was explored by fitting environmental variables to ordination space (in terms of correspondence analysis, CA) with the *envfit* function in R package *vegan* (version 2.5-6) (Oksanen et al. 2019).

### Numeric and statistical analysis, and structural equation modeling

The contribution of different consortia to the total abundance and beta diversity (Bray-Curtis dissimilarity) was calculated with R package *otuSummary* (Yang 2020). A constrained analysis, Canonical Correspondence Analysis (CCA), was used to explore the linkage between microbial community variation and environmental variables (which includes the sediment, pore water chemical composition, and CH_4_ concentration and isotopic signatures) by using R package *vegan* (Oksanen et al. 2019). With the return of CCA, variation partitioning was subsequently performed to resolve the explanatory power of pore water chemistry and sediment nutrient condition by using the *varpart* function in package *vegan*. In addition, structural equation modeling (SEM) was implemented with R package *lavaan* (v0.6.9) (Rosseel 2012) in order to numerically estimate the observed relation between community structure and different environmental features and ecological functionality. Data visualization of environmental data, qPCR and bubble plot was performed with the *ggplot2* package (v3.3.2) (Wickham 2016).

### Data availability

Sequencing data and metadata are deposited at the European Nucleotide Archive (ENA) under BioProject accession number PRJEB49195. The sample accession numbers were listed in Table S1. The metagenomic data was uploaded to ENA under BioProject identifier PRJNA821074.

## Results

### Geochemistry

The pore-water and bulk geochemistry of the two freshwater lakes was very similar, though differences occur in the CH_4_ concentration, CH_4_ carbon isotopes, and iron profiles (Fig. 2). While the results of the lower 300 cm of the lagoon were quite similar to the lakes, large differences in alkalinity, salinity, chloride and sulfate concentrations, pH, and ^13^C-CH_4_ we observed in the upper 300 cm. The C:N ratio was roughly similar over the entire profiles of both lakes, ranging from 7-12. The C:N ratio in samples from 130 - 430 cm in the lagoon is similar to that of the lakes. However, from 0-130 cm and 430 - 510 cm the ratio is substantially higher.

**Fig. 2.**
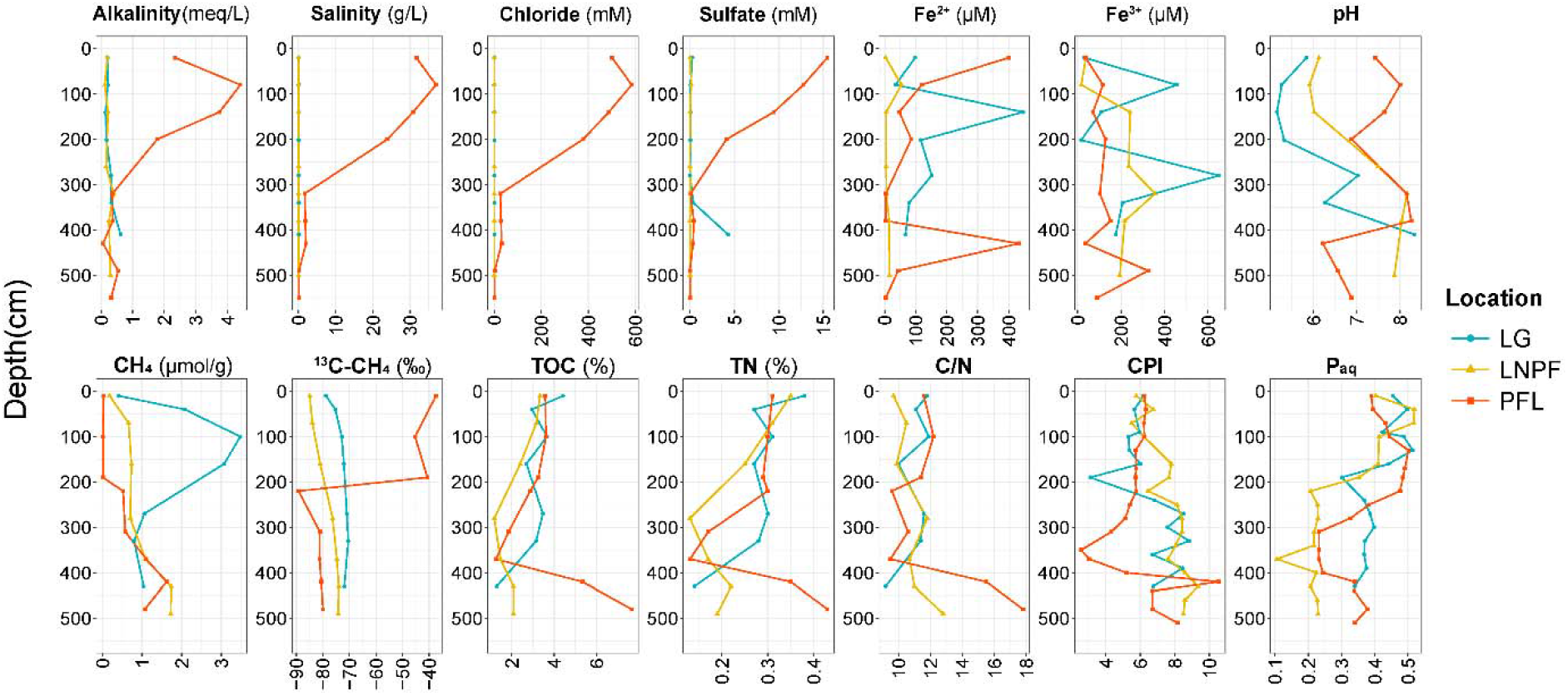
Porewater chemical features of sediments in three cores. The y axis shows the depth from the sediment surface. Water height above sediment surface was 510, 560, and 310 cm for Lake Golzovoye (LG), Northern Polar Fox Lake (LNPF), and Polar Fox Lagoon (PFL), respectively.

### Biomarker Indices

The upper 200 cm of each water body generally had higher P_aq_ values than the rest of the profile, indicating that those layers are composed of more submerged and floating plants, with the deeper sediments having a larger portion of emergent macrophytes and terrestrial plants. The CPI values for both lakes and the lagoon are similar for the top 100 cm, with values between 5.5 - 6.5, converging again at their respective maximum core depths with CPI values of 8-9. In-between these regions the CPI is quite different for each lake indicating differences in the maturity of the OM and decomposition processes, or changes in the vegetation cover. PFL tends to have the most uniform profile over a host of variables, including the biomarker indices.

### Microbial consortia of the methane cycle

Methane cycling microbes which were derived from amplicon sequencing were separated into three main groups: methanogens, aerobic methanotrophs (generally MOBs), and anaerobic methanotrophs (AOMs) (Fig. 3). Across all samples, methanogens were dominated by the methylotrophic Methanomassiliicoccales and Methanofastidiosales, followed by the hydrogenotrophic *Methanoregula* and obligatory acetoclastic *Methanothrix* (commonly referred to as *Methanosaeta*). *Methanosarcina* were scattered in some samples with relatively low abundances.

**Fig. 3.**
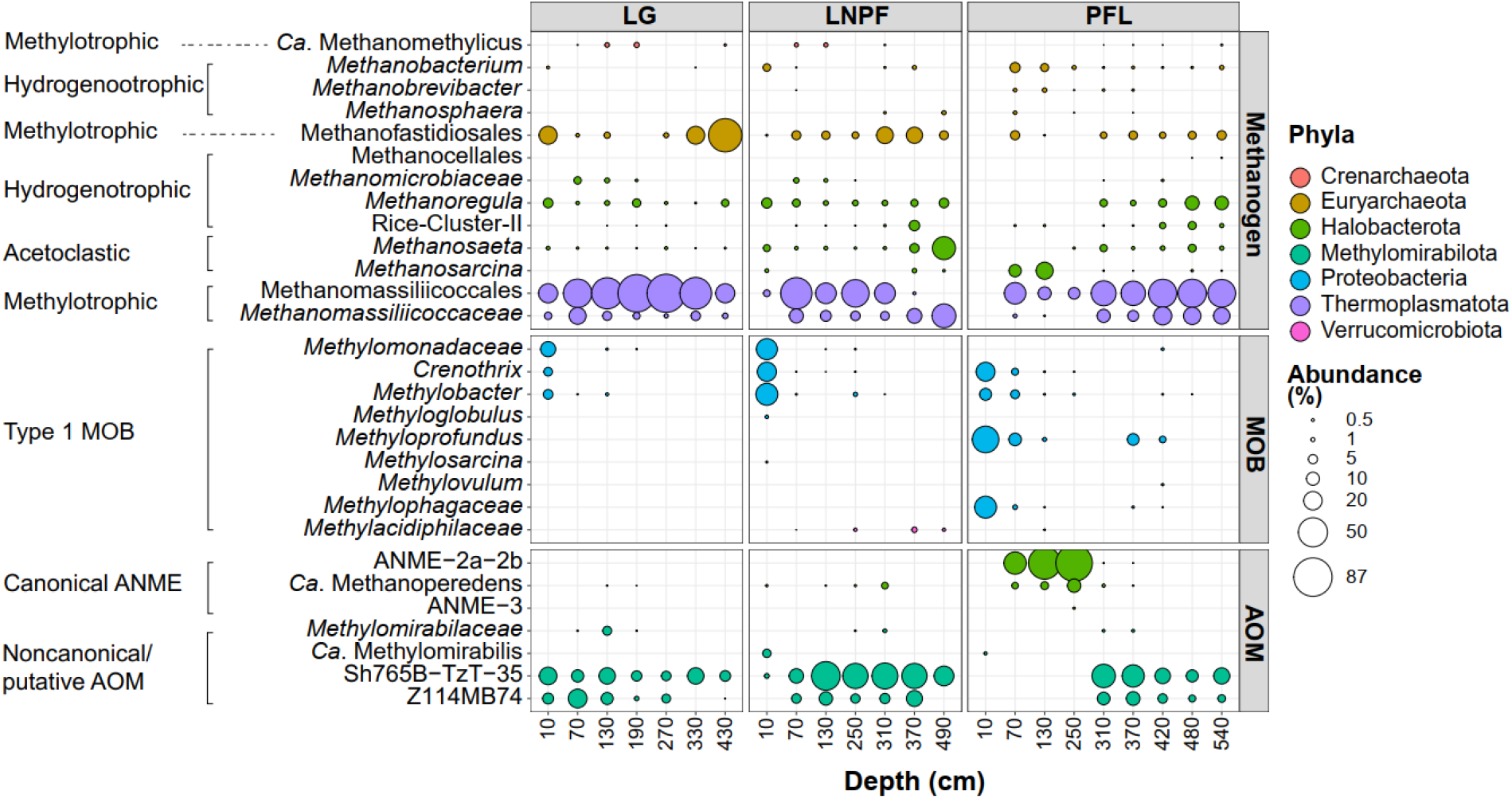
Bubble plot showing the relative abundance within the targeted subset of 264 ASVs (the abundance of all 264 ASVs sum up to 100%). The taxonomy was collapsed at genus level. If an assignment to the genus level was not possible the next higher assignable taxonomic level was used. LG: Lake Golzovoye, LNPF: Northern Polar Fox Lake, and PFL: Polar Fox Lagoon. MOB: methane oxidizing bacteria, AOM: anaerobic methane oxidizers.

Of the anaerobic methanotrophs, ANME-2a/2b is very abundant in the sulfate-rich sediments of the lagoon (upper 300 cm). In addition, there is an abundant group of NC-10 affiliated with potentially non-canonical AOM occurring in the freshwater sediments. Members from Methylomirabiales, especially Sh765B-TzT-35 and Z114MB74, generally prevailed across all samples but were almost absent from the upper 300 cm of the lagoon. Canonical aerobic methane oxidizers (MOBs) have very low abundances except for the upper 10 cm of all three water bodies. Prevalent MOBs include mostly type I Gammaproteobacteria, such as members from *Crenothrix, Methylobacter*, and *Methyloprofundus* which have the highest abundance.

Metagenomic data of PFL sediments demonstrated a high ocurrence of genes responsible for dissimilatory sulfate reduction in the sulfate zone, and the key genes for methanogenesis (*mcrABG*) also exhibited slightly higher abundance in the sulfate zone (Fig. 4). Genes which are involved in dimethylamine and trimethylamine metabolism (*mtbB* and *mttB*) were detected in the sulfate zone while the *mtmB* gene for monomethylamine metabolism was identified beneath the sulfate zone (Fig. 4). The *pmoA* gene which is key for MOB occurred in low abundance in the two upper layers of the sulfate zone. In the sulfate zone of PFL, the *mcrABG* genes was highly associated with the taxonomic lineages of ANME-2a (ANME-2 cluster archaeon HR1, 47.4-59.5%), followed by *Ca*. Methanoperedens (2.7-19.3%) (Table S2). Aligned with the high occurrance of sulfate-dependent ANME2 lineage, abundant sulfate reducing genes and bacteria were also identified in the sulfate zone of PFL.

**Fig. 4.**
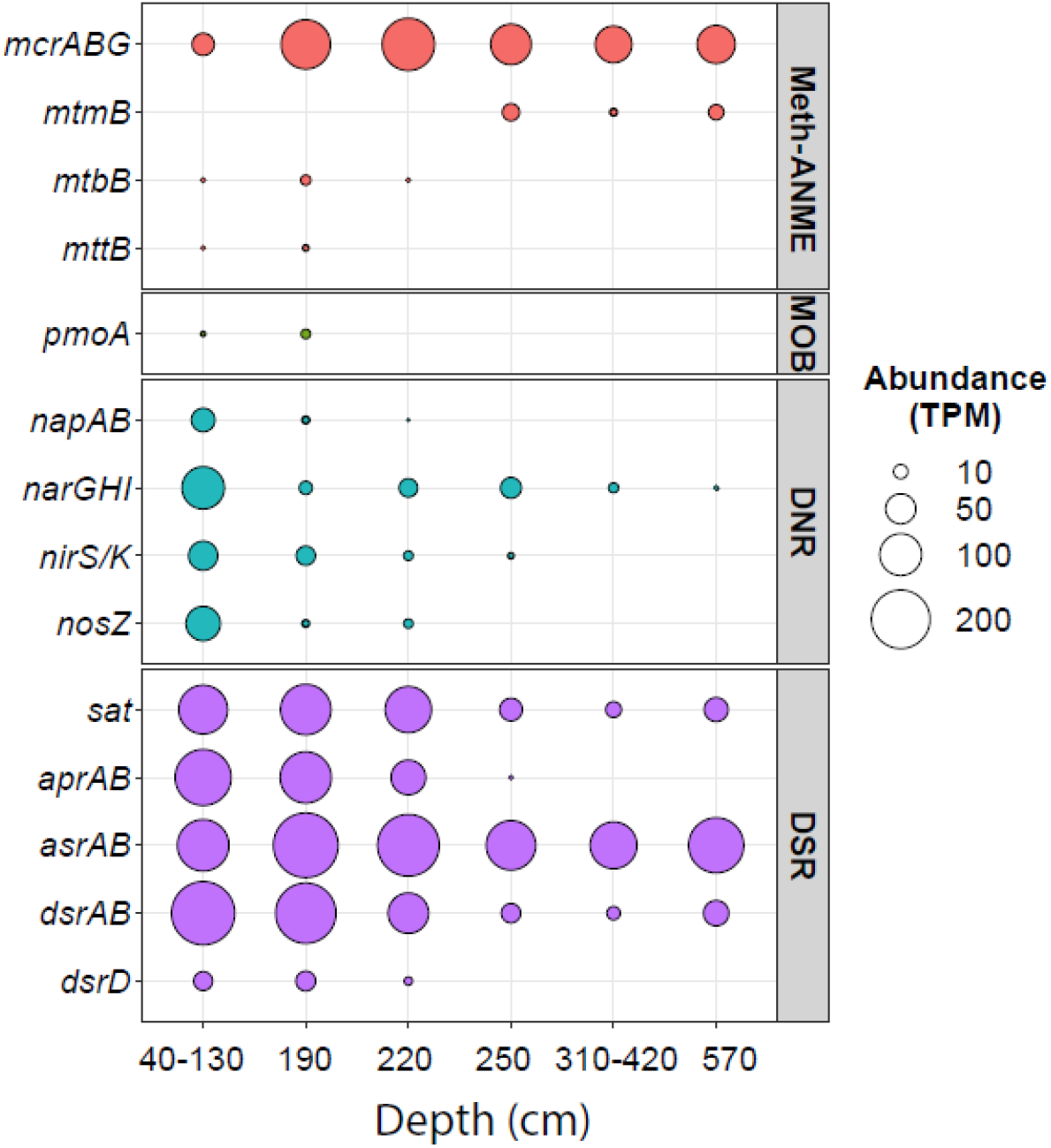
Abundance of key genes involved in methanogenesis/ANME, aerobic methane oxidization (MOB), dissimilatory nitrate reduction (DNR), and dissimilatory sulfate reduction (DSR) over six layers of Polar Fox Lagoon (PFL).

In the ordination space of CCA, the samples from 20-60 cm of PFL are generally separated from the other samples, which represents approximately 74% of the total inertia (i.e. community variations) (Fig. S2a). The variation partition suggested that pore water chemistry was responsible for around 44% of total variance, followed by only 11% contribution by sediment features (Fig. S2b). The SEM further highlights the strong influence of pore water chemistry on microbial assemblages versus the influence of sediment features (Fig. 5). Methanogens and MOB respectively have positive and negative influences on the concentration and isotopic signature of sediment CH_4_ underlining their direct role on the fate of CH_4_.

**Fig. 5.**
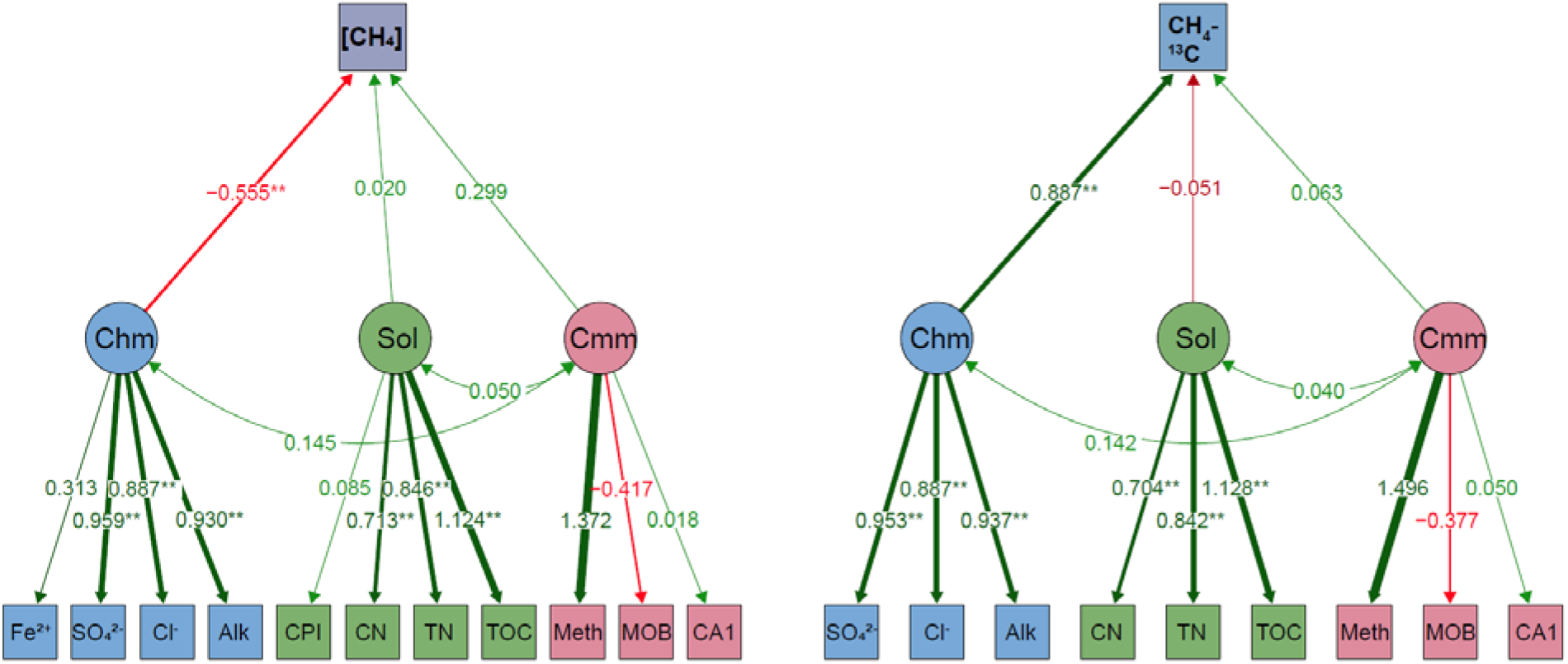
Structural equation modeling showing the influence of the latent variables, pore water chemistry (Chm), carbon and nitrogen (Sol), and Community (Cmm), on the sediment CH_4_ concentration (left) and ^13^C-CH_4_ signature (right). Latent variables are indicated with ovals and observed variables are shown in rectangles. Residual of observed variables were not shown here. Path coefficients were standardized and were significant at *p*-value <0.05, and path width reflect the absolute value of standardized coefficients. The width and color saturation of edges vary with absolute weights. Absolute weights under 0.3 become vaguer in color and smaller in width. Edges with absolute weights under 0.01 are omitted. Meth: methanogen relative abundance, MOB: relative abundance of aerobic methane oxidizing bacteria, CA1: the first axis of corresponding analysis; Alk: alkalinity, [CH_4_]: concentration of sediment methane (µmol g^−1^). The Comparative Fit Indices (CFI) are 0.750 and 0.845 for the left and right analysis, respectively. The Root Mean Square Error of Approximation (RMSEA) are 0.234 and 0.235, respectively. The contribution of AOM had to be in the SEM as the model did not converge due to many zero values of AOM over samples within the limited dataset.

### Co-occurrence and modularity

The quality filtering ended up with 93 ASVs for network construction. These ASVs on average contribute 88.75% (min: 39.38%, first quartile Q_1_: 88.95%, third quartile Q_3_: 96.71%, max: 99.40%) to the total abundance (which constitutes 264 ASVs) and 75.05% (Q_1_: 60.44%, Q_3_: 90.99%) to the total Bray-Curtis dissimilarity (BC) (Table S3). Therefore, the subset of these ASVs largely represent the populations involved in methane cycling.

The species network consists of 93 nodes (ASVs) and 1535 undirected edges (Fig. 6). The density was 0.3588, the average path length was 1.716 edges with a diameter of 3 edges, and the average clustering coefficient transitivity was 0.6848. The centrality measures (degree, betweenness, closeness and eigen centrality) highlighted the importance of the lineages belonging to Methanomassiliicocales, *Methanoregula, Methanofastidiosales* and *Methylomirabilaceae* in the network involved in methane cycling.

**Fig. 6.**
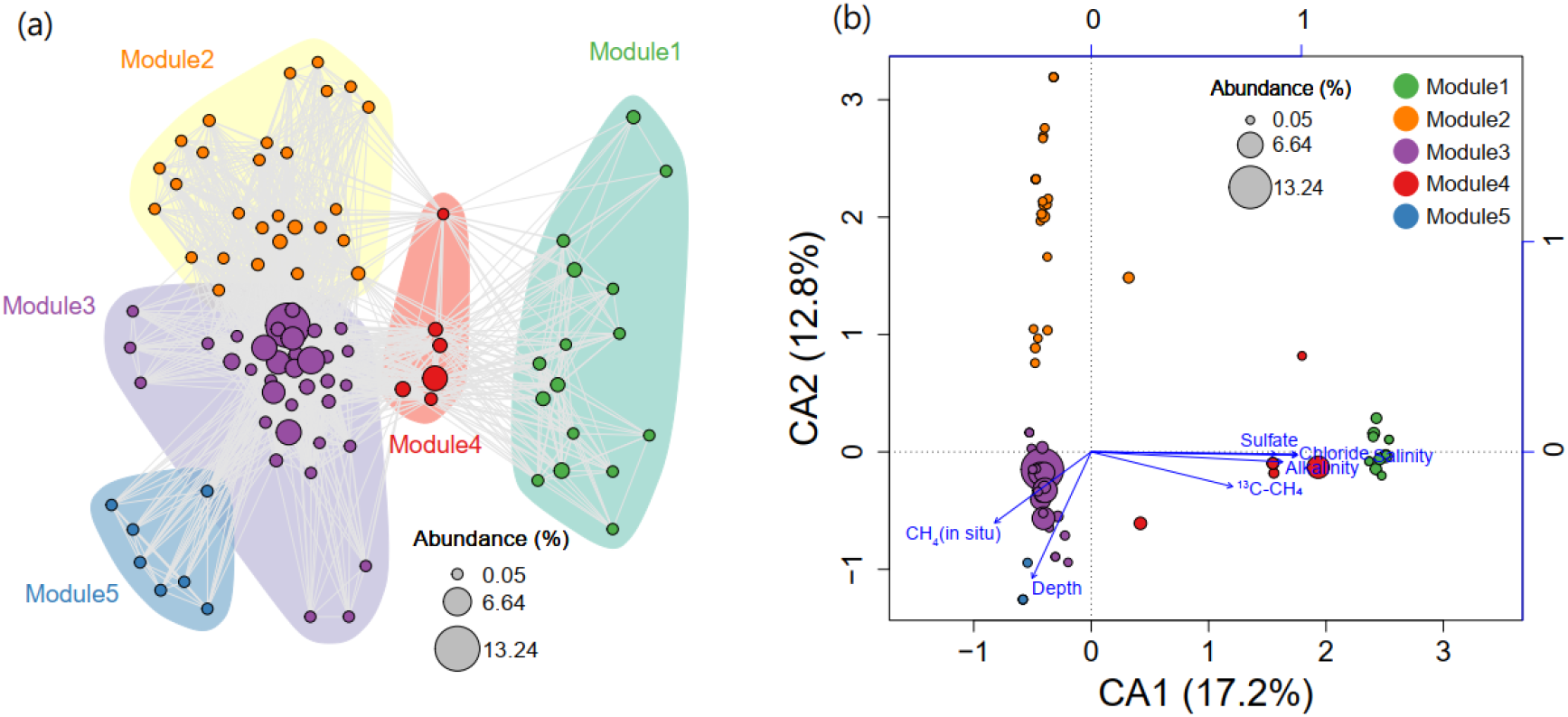
Network modules showing modules for methane cycling consortia (a) and the association of each module and environmental variables and features (b) in correspondence analysis ordination. Bubble size in network and ordination is corresponding to the relative abundance of each ASVs.

Five modules were identified by the *walktrap* algorithm (modularity score =0.204). With the members of each module an ordination shows that Module 1 and 4 are stretched along the first axis, suggesting association with marine-water sulfate, chloride, and alkalinity. In contrast, ASVs in module 2, 3, and 5 are distant from the marine influenced samples, showing stratified cluster features along axis CA2 (Fig. 6 and Fig. S3). The distribution of different modules is likely structured by pore water chemistry. Module 2 occurs mainly in the surface samples (0-10 cm), while module 3 and 5 are distributed in down-core freshwater sediments. Across all modules, keystone taxa consisted of H_2_-dependent methylotrophic methanogens which are mainly affiliated to Methanomassiliicocales or Methanofastidiosales. In module 1 and 4, important components also include methane oxidizers like ANME-2a/b, *Candidatus* Methanoperedens and *Crenothrix*. Every module contains at least one methanogenic lineage.

## Discussion

### Environmental Conditions and History of the Lagoon Sediments

The observed differences in CH_4_ concentrations, isotopes, geochemistry, and microbial assemblages between the lagoon and the two lakes reflect their sediment history and formation. Initial intense thermokarst lake formation on Bykovsky Peninsula began approximately 12 thousand years before present (cal ka BP), before the peninsula itself was shaped (Grosse et al. 2007). While LG evolved independent of the other two water bodies and is younger, PFL and LNPF were originally part of one larger thermokarst basin (Angelopoulos et al. 2020), that went through stages of draining and refilling over several thousand years, eventually becoming and remaining as two separate water bodies within the last 2-4 cal ka BP. The channel between PFL and Tiksi Bay likely formed within the last 2 cal ka BP. Thus, we view LNPF and PFL as two successive stages in the transition of coastal thermokarst landscapes into marine environments. The pore water chemistry of the upper 300 cm of the lagoon sediments indicates that sulfate and other ions have diffused into sediments originally deposited under freshwater conditions, creating a marine-influenced gradient down to approximately 325 cm, below which the porewater chemistry is comparable to the lakes. The mean annual electrical conductivity of Tiksi Bay is 7.1 mS cm^−1^, but values as high as 15 mS cm^−1^ have been observed in winter (Angelopoulos et al. 2020). The seasonal intrusion of water from Tiksi Bay into the lagoon has fostered the formation of a sulfate zone from approximately 0 – 200 cm, followed by a sulfate-methane transition zone (SMTZ) from approximately 200 – 400 cm. This stresses the profound effect that infiltrating water from Tiksi Bay has on the pore-water chemistry of the lagoon sediment, even though the lagoon is only connected to the bay during the ice-free summer months (Angelopoulos et al. 2020). The trends in P_aq_ indicate that overall, OM was of a more terrestrial origin in deeper sediments of all studied water bodies, and has been slowly transitioning towards more submerged and floating plants in the shallower sediments. Though there are some differences in the C:N ratios, all three water bodies stay within a range of 8 – 18 over their entire profile. This further supports that the lagoon was in a similar condition as the freshwater lakes before marine water intrusion. This is also in line with the range found by Schädel et al. (2014) for a variety of permafrost deposits, as well as that found specifically in Yedoma permafrost in the same region (Walz et al. 2018; Wetterich et al. 2008). The C:N ratios could also indicate a large proportion of phytoplankton mixed into the sediments (Biskaborn et al. 2013a; Vyse et al. 2020). The increase in CPI and C:N values in the range of 400 – 500 cm in the lagoon indicates that the OM there is less decomposed compared to the rest of the profile. The less degraded OM is possibly attributed to partially refrozen sediment beneath lagoons water with a cryotic mean annual temperature.

### Lagoon Formation promotes Establishment of AOM Consortia

The CH_4_ distribution and isotopic signatures imply that thermokarst lagoons can establish a strong methane oxidation regime that reduces CH_4_ concentrations compared to their freshwater analogs. Thermokarst lakes in the ice-rich permafrost lowland are generally hotspots of CH_4_ emission (Walter Anthony et al. 2018; Walter et al. 2007), and our results from the sediment columns of LG and LNPF tend to agree with this given CH_4_ concentrations of up to 2.21 µmol g^−1^ at 1.5 m sediment depth in LG. The CH_4_ present in the low sulfate sediment column of all three water bodies is likely a mixture of saturated or supersaturated porewater and small CH_4_ gas bubbles. Calculations under the assumption that all measured CH_4_ was dissolved in the porewater leads to concentrations of up to 17 mM in LG, much higher than the approximately 1.8 - 2.0 mM soluble at 1 bar and 0 - 4°C in freshwater (Pohlman et al. 2013; Yamamoto et al. 1976). In addition, ebullition was observed during sampling in the two freshwater lakes, but not in the lagoon. Spangenberg et al. (2021) and Bussmann et al. (2021) also saw extensive ebullition in LG during their sampling events. Evidence for the presence of CH_4_ bubbles in the sediment or soil column have been observed in other studies in various environments, including marine, lacustrine, and wetland (Pohlman et al. 2013; Thomsen et al. 2001; Tokida et al. 2005). By comparison with the low sulfate sediment columns, the lagoon’s CH_4_ concentration in the sulfate zone was 0.4 – 5% of the rest of the profile and of the concentration along the entire profiles of both lakes. The reduction in porewater CH_4_ concentration is concurrent with heavily enriched δ^13^C-CH_4_ signatures (compared to the lakes), suggesting the occurrence of substantial CH_4_ oxidation. In addition, Spangenberg et al. (2021) found that the CH_4_ concentration in winter ice samples of PFL was as low as 10% of that in LG (PFL mean: 54.7 nM vs LG mean: 645 nM), indicating that CH_4_ was still effectively attenuated in winter when the lagoon is disconnected from the bay. Angelopoulos et al. (2020) found that even though the lagoon was disconnected during the winter, continued growth of the ice cover caused increased salinity in the shrinking water volume during the winter months, likely preserving the marine effect.

There is evidence that methane oxidation in the lagoon is performed by canonical anaerobic methanotrophic archaea, especially the ANME-2a/2b groups. Overall, a profound shift in CH_4_ cycling consortia was observed between the freshwater lakes and PFL, which corresponded to changes in geochemical composition, in particular the high sulfate concentration and salinity. The effect of the geochemical changes was reflected in the environmental partitioning and SEM analysis. Based on the porewater chemistry, such as the Fe^2+^ concentrations, the sediments in all three profiles below the redox boundary, it means roughly at about 10 cm, are likely anoxic (Biskaborn et al. 2013b). Previous studies working on the same lakes have also found that the sediments are anoxic except for limited oxygen availability in the top few centimeters (Jenrich 2020; Schindler 2019; Spangenberg et al. 2021). This would generally inhibit obligate aerobic methane oxidation below this point, providing space for the anaerobic methane oxidizing community to develop, which is supported by the amplicon sequencing and metagenomic results (Fig. 3-4). Accordingly, canonical aerobic MOBs are poorly represented in these thermokarst systems, especially in the zone of PFL where CH_4_ concentrations and isotopes strongly point towards methane oxidation. The metagenomic and amplicon sequencing data support the presence of multiple types of AOMs in the lagoon. The most abundant (Fig. 3) are the S-AOM ANME-2a/2b which are commonly found in classical marine environments. Their dominance is also supported by high sulfate reduction rates in the sulfate zone and SMTZ of the lagoon, which was previously shown to be at least one order of magnitude higher (ca. 50 - 420 nM cm^−3^ d^−1^) than the rest of the profile and the freshwater lakes (Schindler 2019). Strong sulfate reduction in the sulfate zone of PFL could also be supported by the pronounced reduction of sediment CH_4_ concentration and relatively enriched δ^13^C-CH_4_ values (around -40‰) (Fig. 2). Nitrate is also a viable terminal electron acceptor for AOMs. Nitrate concentrations in the pore water, although very low in all three profiles, agree with nitrate concentrations in natural environments, which rarely exceed 50 μM (Schink 2010). Sediment nitrate concentrations of the lagoon being below the detection limit may be a consequence of fast nitrate reduction, rather than a lack of nitrate production. This is supported the by high abundances of genes associated with denitrification and also through occurrence of *Candidatus* Methanoperedens, known to couple AOM with nitrate reduction. Ferric iron (Fe^3+^), which is present in the low sulfate sediments of the lagoon, has also been highlighted as a potential alternative electron acceptor for methane oxidation in many other anoxic habitats (Ettwig et al. 2016; Rooze et al. 2016; Winkel et al. 2018). Such a pathway, according to a recent analysis on eutrophic coastal areas, can play an important role in mitigating CH_4_ by coupling its oxidation with iron, manganese or nitrate reduction beyond the sulfate zone in anoxic sediments due to eutrophication-associated hypoxia (Wallenius et al. 2021). In addition to the results presented here, the work by Schindler (2019) utilizing ^14^C-CH_4_ labelling experiments also found clear evidence of AOM in the 0 – 200 cm lagoon sediments. It is interesting that within the deeper sediments, small pockets of known aerobic MOBs are present, coexisting with canonical AOM and methanogenic organisms, indicating that while this habitat is certainly not ideal for aerobic methanotrophs, it is also not completely inhospitable. Recent studies of other systems found evidence of methane oxidation in anaerobic sediment performed by lineages of canonical aerobic methanotrophs (MOB). For example, in their study of sub-Arctic lake sediments Martinez-Cruz et al. (2017) found that after a six-month incubation under anoxic conditions supporting methane oxidation, type I methanotrophs from the *Methylobacter* genus were the major driver of methane oxidation. This was similar to the findings of Su et al. (2021) in the anoxic sediments of a lake in Switzerland, where type I and type II aerobic methanotrophs were found to grow under both oxic and anoxic incubation conditions. In the lakes and lagoon on Bykovsky peninsula, *Methylobacter*, and other type I methanotrophs are present, however they are most abundant in the top 10-20 cm of sediment and for the most part represent less than 1% of total abundance (Fig. 3). *Crenothrix*, a known aerobic MOB, which could survive under slightly oxygen-deficient conditions in stratified lakes (Oswald et al. 2017), is also concentrated only within the top 10 cm of both lakes and the lagoon similar to other canonical MOB (Fig. 3). Thus, while these organisms may be contributing to the total AOM process in sediments of all three water bodies, their contribution must be considered marginal relative to that of methanotrophic archaea (ANMEs) in the lagoon.

The abundant occurrence of the unclassified environmental groups Sh765B-TzT-35 and Z114MB74 (Fig. 3) closely related with *Candidatus* Methylomirabilales, another known methane oxidizer (Ettwig et al. 2010), imply their ecological importance. The available genomes of Methylomirabilales demonstrated the existence of genes encoding particulate methane monooxygenase (pMMO) (Versantvoort et al. 2018). A recent stable isotope probing (SIP) study in wet forest soil showed that members of Sh765B-TzT-35 act as active anaerobic methanotrophs along with methylotrophic methanogens of *Methanomassiliicoccus*, which were proposed as positive biological indicators of methanotrophy and methanogenesis in wet forest soils (Nakamura 2019). However, our understanding of the role of these clades is extremely poor and it remains open if Sh765B-TzT-35 and Z114MB74 members contribute to methane oxidation in the sediments of this study.

### Lagoon Environments Are Unique from Marine Environments

The combination of molecular and biogeochemical data show that the lagoon forms a microbial habitat unique not only from the freshwater sediments but also from marine sediments. In marine environments, ANME consortia are typically restricted to a SMTZ (Beulig et al. 2019; Egger et al. 2018; Nauhaus et al. 2002), whereas in the lagoon their abundance remains high in the sulfate zone above the observed SMTZ (Fig.3 and Table S2). This could indicate that the location of the SMTZ in the lagoon is seasonally more variable than in marine sediments due to the dynamic nature of the lagoon (e.g. ice buildup and retreat, alternation between freshwater and brackish water inflow) but the reason for the spatially more spread ANME consortia in the lagoon in comparison with marine sediments would require further, more temporally resolved studies.

The observed prevalence of methanogens throughout the sulfate zone and SMTZ would have been a surprising finding under the classical understanding that sulfate reducers outcompete methanogens in high sulfate environments (e.g. Claypool and Kaplan (1974); Sansone and Martens (1981)). However, recent studies have also found co-habitation of methanogenesis and sulfate reduction in a few marine environments (Ozuolmez et al. 2020; Ozuolmez et al. 2015; Sela-Adler et al. 2017), as well as coastal marine sediments (Koebsch et al. 2019; Rooze et al. 2016). This finding is still an important distinction between classical marine environments and a lagoon environment. Mitterer (2010) proposed that the co-occurrence of methanogenesis and sulfate reduction is supported by the presence and utilization of non-competitive methanogenic substrates, such as methylamines and methylated sulfur compounds. Another potential explanation of these differences is the intrusion of freshwater in the lagoon and coastal marine environment. Compared with a lagoon system, marine sediments in the open ocean generally contain a higher sulfate content, while the organic matter is more diluted (Tagliapietra et al. 2009). It is therefore conceivable that marine sediment, aside from coastal regions, also has a reduced amount of available labile carbon. Previous paleoclimate reconstructions have shown that the sediments of the lakes and lagoons on the peninsula are mixtures of permafrost thawed in situ under the waterbody, permafrost eroded from shores by thermo-erosion, and lacustrine sedimentation that has accumulated over approximately 8,000 years (Jongejans et al. 2020). The carbon densities in lagoons in Yedoma permafrost regions are on average 16.5 kg m^−3^, which falls in the range of terrestrial Yedoma (8 kg m^−3^) and thermokarst (24 kg m^−3^) sediments in the Yedoma region (Jenrich 2020; Vyse et al. 2021). The upper three to five meters of sediment contains a higher density of carbon than those below (Jenrich et al. 2021). Moreover, the organic carbon in thermokarst lagoons was strongly correlated with the origin of the organic matter and the depositional conditions (Jenrich et al. 2021; Jongejans et al. 2020). The quality of organic carbon is an important variable regulating microbial carbon turnover (Schuur et al. 2015; Wagner et al. 2005). Likely, thermokarst lakes and lagoons can provide sufficient labile organic carbon for microbial metabolism, in excess of what is usually available in marine environments, and which is another key precondition for the coexistence of sulfate reducing bacteria (SRB) and methanogens (Ozuolmez et al. 2015; Sela-Adler et al. 2017). Therefore, the lagoon provides a sulfate and carbon rich habitat distinctly different from generally carbon poor classical non-coastal marine systems, and based on the prevalence of non-competitive methylotrophic methanogens, may also contain a larger range of organic carbon compounds.

### Methylotrophic Methanogens and the Implications of their Dominance

The methanogenic consortia were dominated by methylotrophic lineages, and their prevalence across all modules and depths (Fig. 3) underscores the ecological importance of methylotrophic methanogenesis in all three water bodies. Methylotrophic methanogenesis has long been considered the least utilized methanogenesis pathway, with many studies instead focusing on hydrogenotrophic and acetoclastic methanogenesis. As Zalman et al. (2018) states, this thinking is so prevalent that rates of acetoclastic methanogenesis from mixed microbial communities have been determined by subtracting the rate of hydrogenotrophic methanogenesis from the total CH_4_ production rate. This method is particularly problematic as methylotrophic methanogens do not appear to be affected by one of the most commonly used acetoclastic methanogenesis inhibitors, methyl fluoride (Penger et al. 2012). As described by Kurth et al. (2020) and Söllinger and Urich (2019), there are a wealth of substrates that various methylotrophic methanogens can and do utilize, many of which are non-competitive. Recent studies have also found that methylotrophic methanogenesis may be the dominant pathway in multiple environments. For example, Xiao et al. (2018) found methylotrophic methanogenesis to be the dominant CH_4_ production pathway in sulfate-rich marine surface sediments from Aarhus Bay in Denmark. Zalman et al. (2018) found that Northern Minnesota peat bogs have a strong potential for methylotrophic methanogenesis based on the amounts of methylated substrate found there. Feldewert et al. (2020) suggested that methylotrophic methanogens are even able to outcompete hydrogenotrophic methanogens for H_2_ when their habitats can provide sufficient methyl groups, even in the presence of sulfate. This study can now be added to the growing list of environments, both freshwater and marine, where methylotrophic methanogenesis is either found to be dominant, or found to have strong potential.

### Niche Differentiation and Stratification of Methane Cycling Consortia in the Lagoon

Changes in biogeochemistry during the transition from freshwater to a lagoon environment promotes unique coexistence and stratification of methane cycling consortia. This is evidenced by the distribution of modules (and therefore microbial consortia) along specific environmental constraints (Fig. 6), as well as the structural equation modeling (Fig. 5). Lacustrine or marine sediments are stratified with respect to available terminal electron acceptors (TEAs), nutrients, and redox potential, as well as microbial consortia. The niche differentiation and stratification in PFL versus the two lakes is conceptualized in Figure 7. Accordingly, the formation of the sulfate zone during lagoon development likely took a prominent role in niche separation. As a consequence, the microbial consortia in different zones and niches were reconstructed, and these new species interactions lead to the coexistence of consortia which were fit for these new microhabitats in the lagoon. Examples for this are the predominance of ANME 2a/2b and *Ca*. Methanoperedens in the sulfate zone of PFL and the co-occurrence of certain groups of *Methanomassiliicoccaceae*, which have a high affinity or preference for marine-water influenced conditions (Fig. 3). The importance of niche separation in regulating the stratified distribution and separation of microbial community has been shown before in various habitats, especially in sediments (Grunke et al. 2011; Marchand et al. 2004; Nunoura et al. 2016; Vuillemin et al. 2018). Therefore, the associated environmental controls on different modules highlights the importance of niche differentiation and environmental heterogeneity between the lakes and the lagoon. At the same time, the network analysis suggests that methylotrophic methanogens constitute generalists in thermokarst lake and lagoon sediments.

**Fig. 7.**
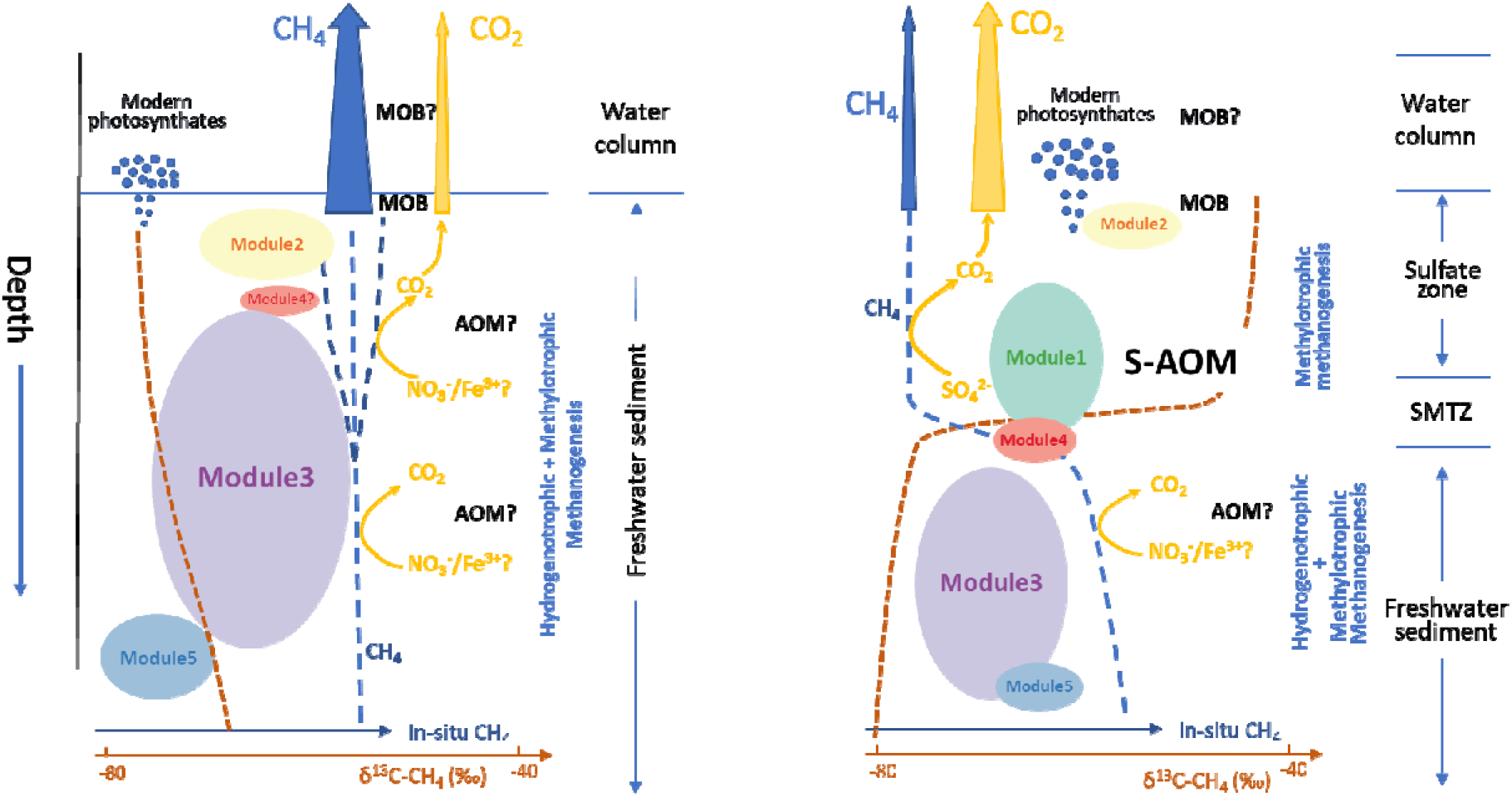
Schematic plot showing the stratified distribution of methane-cycling communities, processes in freshwater thermokarst lakes (left) and re-assembly of community and methane turnover pathways after thermokarst lakes turn into thermokarst lagoons (right).

## Conclusion and Perspectives

In Arctic coastal permafrost regions, the transition from thermokarst lakes to lagoons changes the ionic strength and oxidative capacity, and introduces new microbial species. Consequently, the initial geochemical and trophic stratification is reorganized which in turn induces profound shifts in the structure and functionality of the methane-cycling community, and influences carbon feedbacks in the lagoon by supporting a strong AOM community. The results of our study therefore indicate that, while considerably different from classical marine environments, lagoon environments can still be favorable habitats for canonical AOM consortia. In the lagoon system, a high load of organic carbon and non-competitive substrates further afford the co-existence of methanogens and other sulfate-reducing organisms. Owing to effective methane oxidation, thermokarst lagoons could play an indispensable role in mitigating methane release from thermokarst ecosystems. An estimation of the current extent of pan-Arctic lagoons including lagoons in the Teshekpuk Lake region, Baldwin Peninsula, Mackenzie Delta, Taimyr Peninsula, Lena Delta, and Tiksi regions, which is detailed in the supplement, shows that these lagoons currently occupy an estimated area of 2579 km^2^, which is roughly equivalent to the size of the country of Luxemburg. Although still comparatively small in area, thermokarst lagoons transform microbial methane cycling communities and will further expand in Arctic regions, especially along the Laptev Sea coast due to sea-level rise, accelerated permafrost thaw, intensified coastal erosion, and changing sea ice regimes. Pan-arctic lagoons may therefore be more relevant to the permafrost carbon budget than is currently reflected in the literature, especially if they host a strong AOM community like shown here. Beyond that, thermokarst lagoon systems represent important natural laboratories for studying the effect of sea level rise on vulnerable, organic-rich coastal permafrost landscapes. Therefore, their potential to serve as habitat for efficient AOM-communities deserves more attention in the future.

## Acknowledgements

We thank Jan Kahl, Lutz Schirrmeister, and Axel Kitte for assisting with drilling during field work. We thank Daniela Warok for help with lipid biomarker analysis. We also thank Anke Saborowski for her contribution to the qPCR analysis. This study was supported by the Helmholtz Gemeinschaft (HGF) through funding for SL’s Helmholtz Young Investigators Group [VH-NG-919]. SY and SL were supported by the German Ministry of Education and Research as part of the projects CarboPerm [grant no. 03G0836A, 03G0836D], and KoPf [grant no. 03F0764A, 03F0764F]. SY also acknowledges support from Chinese Academy of Sciences. GG and JS were supported by ERC PETA-CARB (#338335) and the HGF Impulse and Networking Fund (#ERC-0013). MJ was supported by DBU grant (project “Characterisation of organic carbon and estimation of greenhouse gas emissions in a warming Arctic”). Field work was supported by funds from AWI and GFZ as well as the ERC PETA-CARB project.

