## Supplemental Information for "Anaerobic methane oxidizing archaea offset sediment methane concentrations in Arctic thermokarst lagoons"

**Supplementary material**

**Quantitative PCR assays**

Quantitative PCR (qPCR) was performed to estimate the abundance of total community and specific consortia by using universal primer (Eub341-F/Eub534-R), *mcrA* primer (mlas/mcrA-rev), *pmoA* (189F/661R), *dsrB* (dsrB2060-F/dsrB4-R) and the specific primer set for *Candidatus Methanoperedens* (Vaksmaa et al. 2017) according to Unger et al. (2021) and Winkel et al. (2018). The qPCR was runs in technical triplicates on a CFX96 real-time thermal cycler (Bio-Rad Laboratories Inc., USA), with amplification efficiencies ranging from 91 to 99%. The qPCR results were normalized by the amount of DNA, in ng extracted per gram of fresh material.

**Gene copy numbers**

The qPCR results (Fig. S4) show that the lagoon had, for most depths, more 16S rRNA gene copies than either of the lakes, suggesting a higher microbial abundance in the lagoon. The lagoon also had more copies of the *pmoA* gene than both GL and PFL, indicating a larger population of *pmoA*-bearing methanotrophs over the entire profile. Interestingly the copies of the *mcrA* gene are roughly equivalent in all three water bodies, indicating that the size of the methanogenic community is roughly the same for each. *Ca.* Methanoperedens and *dsrB* gene sequence copies are slightly more abundant in the upper 200 cm of the lagoon than in the lakes, and roughly equivalent in the lower half of the profile. This suggests that there is a difference in the methanotroph population in the lagoon in general, and a difference in a larger portion of the microbial community between the upper 200 cm of the lagoon and the lakes.

**Table S1**. Sample accession numbers in the European Nucleotide Archive used in this study. Sample ID’s PG2420, PG2423, and PG2426 refer to LG, PFL, and LNPF, respectively. LG: Lake Golzovoye; PFL: Polar Fox Lagoon; LNPF: Lake North Polar Fox. The last letter ‘a’ and ‘b’ in the sample identifiers denotes the replicates of each sample.

| Sample aliens | Sample ID | Accession | Sample ID | Accession |
| --- | --- | --- | --- | --- |
| LG_10 | PG2420_10a | ERS8483289 | PG2420_10b | ERS8483290 |
| LG_70 | PG2420_70a | ERS8483293 | PG2420_70b | ERS8483294 |
| LG_130 | PG2420_130a | ERS8483297 | PG2420_130b | ERS8483298 |
| LG_190 | PG2420_190a | ERS8483301 | PG2420_190b | ERS8483302 |
| LG_270 | PG2420_270a | ERS8483305 | - | - |
| LG_330 | PG2420_330a | ERS8483308 | PG2420_330b | ERS8483309 |
| LG_430 | PG2420_430a | ERS8483312 | PG2420_430b | ERS8483313 |
| PFL_10 | PG2423_10a | ERS8483314 | PG2423_10b | ERS8483315 |
| PFL_70 | PG2423_70a | ERS8483318 | PG2423_70b | ERS8483319 |
| PFL_130 | PG2423_130a | ERS8483322 | PG2423_130b | ERS8483323 |
| PFL_250 | PG2423_250a | ERS8483330 | PG2423_250b | ERS8483331 |
| PFL_310 | PG2423_310a | ERS8483334 | PG2423_310b | ERS8483335 |
| PFL_370 | PG2423_370a | ERS8483338 | PG2423_370b | ERS8483339 |
| PFL_420 | PG2423_420a | ERS8483342 | PG2423_420b | ERS8483343 |
| PFL_480 | PG2423_480a | ERS8483348 | PG2423_480b | ERS8483349 |
| PFL_540 | PG2423_540a | ERS8483350 | PG2423_540b | ERS8483351 |
| LNPF_10 | PG2426_10a | ERS8483352 | PG2426_10b | ERS8483353 |
| LNPF_70 | PG2426_70a | ERS8483356 | PG2426_70b | ERS8483357 |
| LNPF_130 | PG2426_130a | ERS8483360 | PG2426_130b | ERS8483361 |
| LNPF_250 | PG2426_250a | ERS8483368 | PG2426_250b | ERS8483369 |
| LNPF_310 | PG2426_310a | ERS8483372 | PG2426_310b | ERS8483373 |
| LNPF_370 | PG2426_370a | ERS8483376 | PG2426_370b | ERS8483377 |
| LNPF_490 | PG2426_490a | ERS8483384 | PG2426_490b | ERS8483385 |

**Table S2**, Taxonomic composition (%) showing lineages which contain *mcrABG* genes based on the metagenomic data for Polar Fox Lagoon. Depth of PFL1: 40-130 cm, PFL2: 190 cm, PFL3: 220 cm, PFL4: 250 cm, PFL5: 310- 480 cm, PFL6: 570 cm

| Taxonomy | PFL1 | PFL2 | PFL3 | PFL4 | PFL5 | PFL6 |
| --- | --- | --- | --- | --- | --- | --- |
| ANME-2a cluster archaeon HR1 | 59.46 | 41.33 | 47.37 | 0 | 0 | 0 |
| *Ca*. Methanoperedens | 2.70 | 19.93 | 0 | 0 | 0 | 0 |
| *Methanoregula* | 0 | 0.37 | 21.05 | 87.50 | 42.86 | 56.48 |
| Methanomicrobiales | 5.41 | 0.37 | 10.53 | 12.50 | 57.14 | 43.52 |
| Euryarchaeota | 32.43 | 38.01 | 21.05 | 0 | 0 | 0 |

**Table S3**: Contribution of 93 ASVs to total abundance and Bray-Curtis dissimilarity. Please note the mean value of relative abundance lower than the 1^st^ quantile, that is because of the left-skewed data which pulled down the mean values.

|  | Min. | 1st quartile | mean | 3rd quartile | Max. |
| --- | --- | --- | --- | --- | --- |
| Relative abundance (%) | 39.38 | 88.95 | 88.75 | 96.71 | 99.4 |
| Bray-Curtis dissimilarity | 0.2062 | 0.6044 | 0.7505 | 0.9099 | 0.9866 |

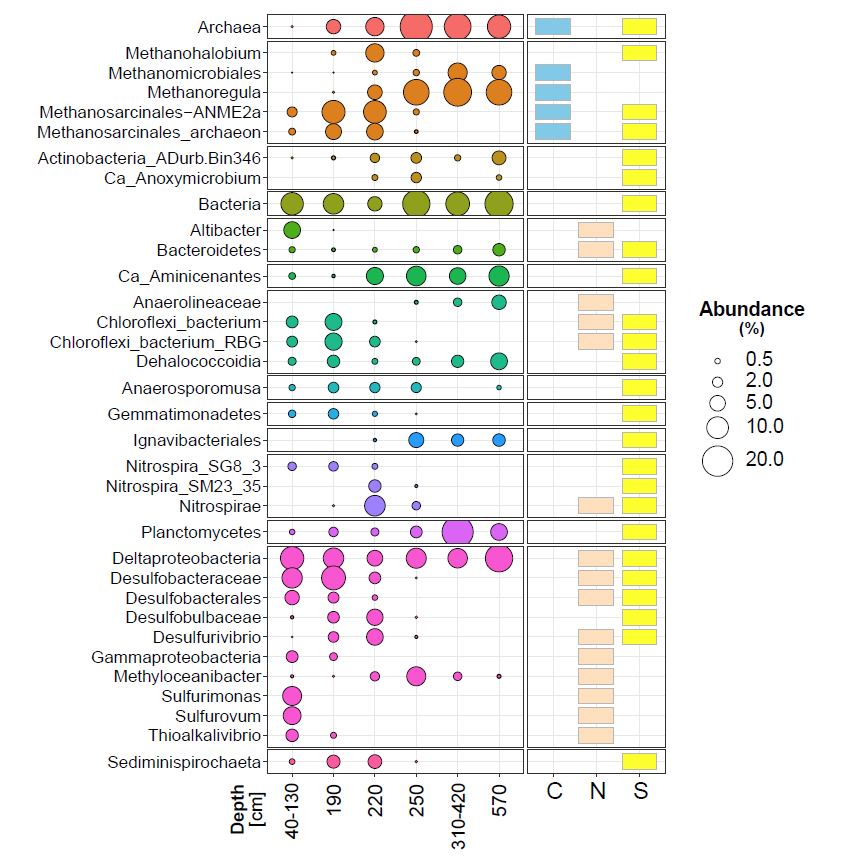

**Fig. S1**, Compositional variation of taxa annotated from the key genes from Fig. 3 for 6 layers of the Polar Fox Lagoon (PFL). The right panel demonstrates whether the presence/absence of key genes involved in C (methanogenesis), N (dissimilatory nitrate reduction) and S (dissimilatory sulfate reduction). The clearest taxonomy was given for each lineage. The taxa with mean relative abundance >0.5% over samples were shown in this plot.

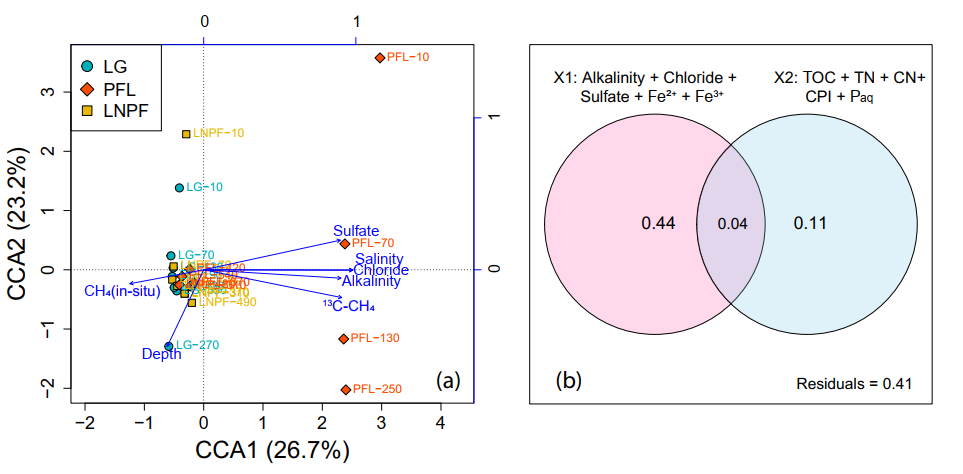

**Fig. S2.** Variation partitioning on constrained correspondence analysis showing the explanation power of two categories of environmental features (X1: pore water chemistry and X2: sediment nutrient condition).

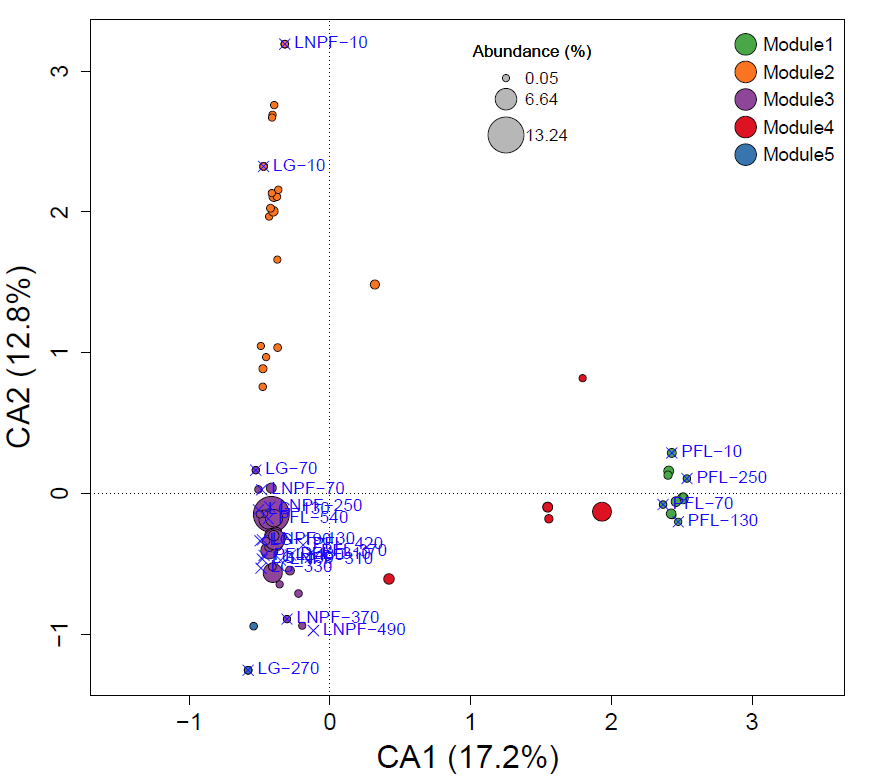

**Fig. S3**, Correspondence Analysis (CA) showing association between samples and ASVs which were used in the network analysis. The points denote individual putative species (ASVs) which were colored by module membership. The blue crosses show different samples in the ordination space. This is a supplementary plot to Figure 6b in the main text which did not show samples and was used to highlight the linkage between module members and environmental variables, while here environmental items were not shown to show the prevalence of module members overs samples. According to the analysis, module 1 and module 4 are more representative in the sulfate-zone and SMTZ of PFL, module 2 occur more in the upper layers of freshwater thermokarst sediments, module 3 and 5 prevail in the lower, non-saline (fresh water) layers across all cores in this study.

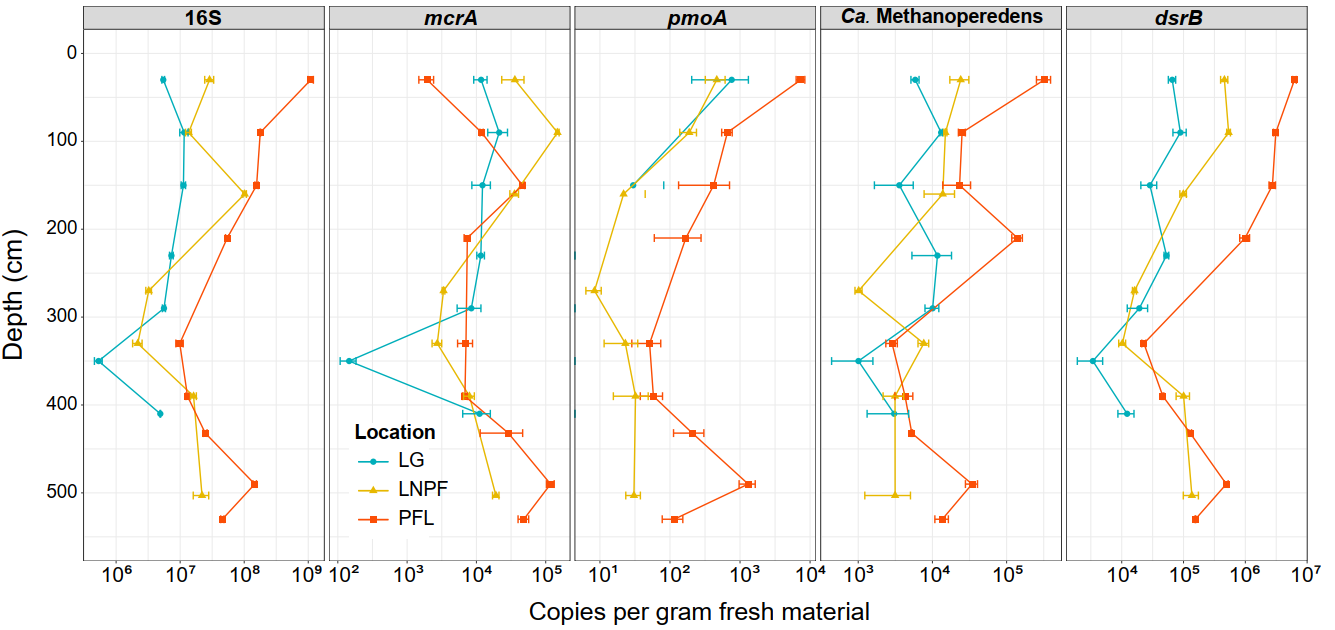

**Figure S4**. Quantitative PCR showing the gene copies of 16S, *mcrA*, *pmoA*, *dsrB* and the *Candidatus* *Methanoperedens*. Gene copies were normalized by gram of fresh material used for DNA extraction.
